# Formate reduces ischemic injury in female hearts lacking alcohol dehydrogenase 5

**DOI:** 10.1101/2025.07.29.667470

**Authors:** Haley Garbus-Grant, Obialunanma V. Ebenebe-Kasonde, Raihan Kabir, Mark J. Kohr

## Abstract

Ischemic heart disease is a primary cause of death for men and women in the United States. Recent epidemiologic findings, however, suggest that pre-menopausal women have inherent protection from many cardiovascular pathologies compared to age-matched men, which is lost with menopause. We and others have documented similar protective signaling in animal models, with females exhibiting protection from ischemic injury that is lost with ovariectomy (OVX). Furthermore, in recent studies, we demonstrated that the loss of alcohol dehydrogenase 5 (ADH5) blocked sex-specific cardioprotection in females, but activation of aldehyde dehydrogenase 2 (ALDH2) provided a rescue. ADH5 and ALDH2 both metabolize formaldehyde to formate, potentially implicating formate in female-specific cardioprotection. Therefore, the objective of this study was to examine a role for formate during ischemic injury in female hearts using wild-type (WT) and ADH5^-/-^ mice. We also aimed to explore estrogen-dependent effects by using ovariectomized (OVX) WT mice. To assess the protective effects of formate in intact female WT and ADH5^-/-^ hearts, as well as OVX WT hearts, hearts were Langendorff-perfused and subjected to ischemia/reperfusion (I/R) injury. Since formate is used in one-carbon metabolism (OCM), select OCM enzymes were also probed via western blot. Importantly, we found that formate significantly reduced infarct size in female ADH5^-/-^ hearts subjected to I/R injury, but formate was without effect in intact female WT hearts. Additionally, formate failed to reduce I/R injury in OVX WT hearts, despite OVX WT hearts exhibiting reduced ADH5 and ALDH2 activity. However, we noted that the expression of certain OCM enzymes was downregulated in OVX WT hearts vs. intact WT females, which may prevent proper formate utilization by OCM in OVX WT hearts. Furthermore, blockage of formate import into OCM in intact female WT hearts also exacerbated I/R injury. Taken together, our findings support formate utilization by OCM as a key component of cardioprotective signaling in female hearts, with estrogen acting as a potential mediator.

## INTRODUCTION

Ischemic heart disease (IHD) is a major cause of death among men and women in the United States.^1^ Epidemiologic studies, however, suggest that premenopausal women are at reduced risk for developing IHD compared to age-matched men, but IHD incidence sharply increases with the menopausal transition.^2–7^ Female-specific protective mechanisms have also been described by our group and others in animal models of ischemia-reperfusion (I/R) injury, with protection disrupted via ovariectomy (OVX),^8–17^ supporting a cardioprotective role for estrogen in female hearts. Unfortunately, hormone replacement therapy has largely failed to block the progression of CVD in postmenopausal women.^18, 19^ As such, our understanding of the relationship between estrogen and female-specific cardioprotective signaling is incomplete.

Our prior work demonstrated that female hearts exhibit enhanced endothelial nitric oxide synthase (eNOS) expression and activity, nitric oxide (NO) production, and protein *S*-nitrosation,^8–10^ all of which are thought to contribute to female-specific protection from ischemic injury. Female hearts also show increased activity of alcohol dehydrogenase 5 (ADH5),^8–10^ an enzyme that regulates protein *S*-nitrosation levels and metabolizes formaldehyde to formate. More recently, we found that the inhibition or knockout of ADH5 increased post-ischemic formaldehyde levels and exacerbated ischemic heart injury in females, but ADH5^-/-^ hearts could be rescued with the activation of aldehyde dehydrogenase (ALDH2).^10^ As both ADH5 and ALDH2 metabolize formaldehyde to formate, our findings suggest these enzymes contribute to cardioprotection by either metabolizing formaldehyde or generating formate. Prior work has shown that mice lacking both ADH5 and ALDH2 have reduced lifespans,^20, 21^ likely because of formaldehyde-induced toxicity. Conversely, formate can also be used in one-carbon metabolism (OCM),^22^ which is a critical process for generating important biomolecules, including antioxidants, that are critical for combating I/R injury. However, the role of ADH5, formate and OCM remain relatively unexplored in the female heart.

In the current study, we investigated a protective role for formate in ischemic injury using female wild-type (WT) and ADH5^-/-^ hearts. We also examined a role for estrogen in the regulation of ADH5 and ALDH2 activity using a model of ovariectomy (OVX) and further investigated the potential for formate-induced protection in hearts from OVX WT mice. While formate provided little benefit to female WT hearts, hearts from ADH5^-/-^ females showed a significant reduction in infarct size with exogenous formate perfusion. We also found reduced ADH5 and ALDH2 activity in OVX WT hearts, but despite this decrease in activity, exogenous formate failed to reduce ischemic injury in OVX WT hearts. Importantly, inhibition of formate intake into the OCM pathway exacerbated I/R injury in WT female hearts. Taken together, our findings support endogenous and exogenous formate utilization by OCM as a key component of cardioprotective signaling in female hearts. Moreover, estrogen may act as a potential mediator of this female-specific protective signaling pathway.

## MATERIALS AND METHODS

### Animals

This investigation conforms to the Guide for the Care and Use of Laboratory Animals published by the National Institutes of Health (NIH; Publication No. 85-23, Revised 2011) and was approved by the Institutional Animal Care and Use Committee of the Johns Hopkins University. Intact and ovariectomized (OVX) female mice were purchased from the Jackson Laboratory (Bar Harbor, ME) and used between 15-30 weeks of age. OVX mice underwent ovariectomy at 8 weeks of age at the Jackson Laboratory and were given three weeks for recovery and estrogen clearance before shipment. Generation of ADH5^-/-^ mice was described previously;^10^ ADH5^-/-^ mice were used between 15-30 weeks of age. Mice were housed in a pathogen-free environment, maintained on standard lab chow, and acclimated to our animal facility for at least one week before experiments.

### Langendorff Heart Perfusion

Mice were anesthetized with a mixture of ketamine (90 mg/kg; Hofspira) and xylazine (10 mg/kg; Sigma-Aldrich) via intraperitoneal injection and were anticoagulated with heparin (Fresenvis Kabi, Lake Zurich, IL). Hearts were excised and canulated on a Langendorff apparatus, and perfused retrogradely with Krebs-Henseleit buffer (KHB) (95% O_2_, 5% CO_2_, pH 7.4) under constant pressure (100 cmH_2_O) and temperature (37°C) as previously described.^8–10, 23–25^ KHB consisted of (in mmol/L): NaCl (120), KCl (4.7), KH_2_PO_4_ (1.2), NaHCO_3_ (25), MgSO_4_ (1.2), d-glucose (11), and CaCl_2_ (1.75). Hearts undergoing molecular analysis were perfused for 5 minutes and snap-frozen in liquid nitrogen, while hearts undergoing I/R injury were subjected to the protocol described below.

### Ischemia/Reperfusion Injury Protocol

Hearts were subjected to I/R injury as previously described.^8–10, 23–25^ Briefly, hearts were equilibrated on the Langendorff apparatus for 10 minutes with KHB and then perfused with or without 30 μmol/L formate (Sigma-Aldrich, St. Louis, MO) in KHB for 20 minutes. The flow of buffer was then stopped for 25 minutes to induce global, normothermic ischemia. After ischemia, the flow of buffer was restored, and hearts were re-perfused for 90 minutes in KHB with or without 30 μmol/L formate. This protocol was performed in the dark for all experimental groups. Finally, hearts were perfused and incubated with tetrazolium trichloride (TTC, Sigma-Aldrich) dissolved in KHB for 20 mins at 37°C, and placed in formalin for fixation. After formalin fixation for a minimum of 24 hours, hearts were cut into 1-2 mm cross-sections and imaged using a dissecting scope (Leica). Infarct size was then assessed using ImageJ software (NIH, Bethesda, MD) and expressed as a percentage of the total area of the ventricle as previously described.^8–10, 23–25^ All samples were blinded prior to sectioning and infarct analysis.

### Tissue Homogenization

Snap-frozen hearts were homogenized in 1x cell lysis buffer (Cell Signaling Technology, Danvers, MA) supplemented with a protease/phosphatase inhibitor cocktail (Cell Signaling Technology) and neocuproine (Sigma-Aldrich) unless otherwise noted. Mechanical dissociation was performed using a Precellys Evolution 24 homogenizer (2x30s cycles, 0°C, 7200 RPM; Bertin Instruments, Montigny-le-Bretonneux, France) using a hard tissue lysing kit. Homogenates were assayed for protein concentration via Bradford assay, and aliquots of tissue homogenates were stored at 80°C.

### Aldehyde Dehydrogenase Activity Assay

Snap-frozen hearts were homogenized in Dulbeco’s phosphate-buffered saline (Gibco, Carlsbad, CA) with a protease/phosphatase inhibitor cocktail (Cell Signaling Technologies) and neocuproine. After a 20-minute incubation on ice, samples were centrifuged at 16,000 g at 4°C for 20 minutes. Protein samples (630 μg) were then diluted in assay buffer and ALDH2 activity was assessed using a commercial colorimetric kit (ab115348; Abcam, Cambridge, MA) per the manufacturer’s instruction. Activity was estimated by measuring the conversion of NAD^+^ to NADH via absorbance at 450 nm every minute for 120 minutes at 25°C.

### Alcohol Dehydrogenase 5 Activity Assay

ADH5 activity was assessed in whole heart homogenates as previously described,^8–10^ in the dark. Whole heart homogenates (100 μg) were diluted in assay buffer containing (in mmol/L): Tris-HCl pH 8.0 (20), EDTA (0.5), neocuproine (0.5) with 0.1% NP-40 and protease/phosphatase inhibitor (Cell Signaling). NADH (200 mmol/L, Sigma-Aldrich) and GSNO (400 mmol/L, Sigma-Aldrich) were then added to initiate the reaction. Activity was estimated by measuring the consumption of NADH via absorbance at 340 nm every minute for 30 minutes at 25°C.

### Formaldehyde Assay

Formaldehyde levels were assessed in whole heart homogenates from intact and OVX WT hearts using a commercial kit (Sigma-Aldrich) as previously described.^10^ Whole heart homogenates (800 μg) were initially deproteinated with trichloroacetic acid and neutralized prior to the assay, and the resulting supernatant was diluted in assay buffer. Formaldehyde levels were measured via fluorescence, and a standard formaldehyde curve was used to determine the concentration.

### Western Blot

Whole heart homogenates (30 μg) were separated (20 mins at 75 V; 70 mins at 175 V) on a gradient Bis-Tris SDS-PAGE gel (4-12%, NuPAGE; Thermo Fisher, Carlsbad, CA). Proteins were transferred (90 mins, 220 mA, 30V) to a PVDF membrane (Thermo Fisher) and total protein was quantified using Ponceau S stain (Thermo Fisher). Membranes were then blocked in (5% wt/vol) bovine serum albumin (Sigma-Aldrich) for 1 hour at room temperature or overnight at 4°C, followed by incubation with respective primary antibodies overnight at 4°C in blocking solution: methionine synthase (1:1000, Rabbit mAb, Cell Signaling #68796), lysine-specific histone demethylase 1 (1:1000, Rabbit mAb, Cell Signaling #2184), methylenetetrahydrofolate dehydrogenase 1 like (1:1333, Rabbit mAb, Cell Signaling #14999), methylenetetrahydrofolate dehydrogenase 2 (1:1000, Rabbit mAb, Abcam ab228757), methyltetrahydrofolate reductase (1:1000, Rabbit mAb, Abcam ab203786), ALDH2 (1:1000, Rabbit mAb, Cell Signaling, #18818), serine hydroxymethyltransferase 1 (1:1000, Rabbit Ab, Cell Signaling #80715) and 2 (1:1000, Rabbit Ab, Cell Signaling #12762), and thymidylate synthase (1:1000, Rabbit Ab, Cell Signaling #9045). Membranes were washed with 1x Tris-buffered saline with 0.1% Tween (TBS-T) at room temperature and then incubated for 1 hour with anti-rabbit HRP-linked secondary antibody (1:5000, Cell Signaling #7074) at room temperature. Membranes were again washed in TBS-T. Chemiluminescence substrate (SuperSignal West Pico PLUS, Thermo Fisher) was added to the membrane and visualized using an iBright imager (Thermo Fisher). Protein expression was quantified using ImageJ software (NIH).

### Statistical Analysis

Data were analyzed using GraphPad Prism (La Jolla, CA). Results are expressed as the mean +/- standard error of the mean. Outliers were identified using the ROUT method (Q = 1%), and normality of distributions was assessed using a Shapiro-Wilk test, as described previously. Significance was determined using a Students t-test or a one-way ANOVA with Tukey’s multiple comparisons for normally-distributed data. For non-normally distributed data, significance was determined using the Mann-Whitney rank sums test for two groups or a Kruskal-Wallis test with Dunn’s multiple comparisons correction. Significance was set at p<0.05. For enzyme assays, the linearity of the data was determined by a best-fit line. Linear regression analysis was used for statistical comparison of linear data, (i.e. enzyme activity assays). Curve differences were evaluated with 95% confidence intervals.

## RESULTS

### Formate rescues female ADH5-/- hearts from ischemic injury

Our prior work demonstrated that the loss of ADH5 increased post-ischemic formaldehyde levels and exacerbated I/R injury in female hearts.^10^ Interestingly, ALDH2 activation provided a rescue. These findings suggest that formaldehyde toxicity or the loss of formate production contributes to ischemic injury in female hearts lacking ADH5. In the current study, we tested the latter hypothesis. WT and ADH5^-/-^ female hearts were Langendorff-perfused with or without 30 μmol/L formate for 20 minutes, subjected to 25 minutes of ischemia, and then re-perfused with or without 30 μmol/L formate for 90 minutes. Formate perfusion significantly reduced infarct size in female ADH5^-/-^ hearts, providing a rescue for the loss of ADH5 (Fig 1A-B). However, formate did not confer a protective benefit in WT female hearts (Fig. 1A-B). Surprisingly, male ADH5^-/-^ hearts, which we have already demonstrated to be protected from ischemic injury,^10^ showed a further reduction in infarct size with formate perfusion (Fig. 1A-B). These findings demonstrate that formate yields protection from I/R injury in the absence of ADH5 and further support a role for formate in female-specific cardioprotection.

**Figure 1.**
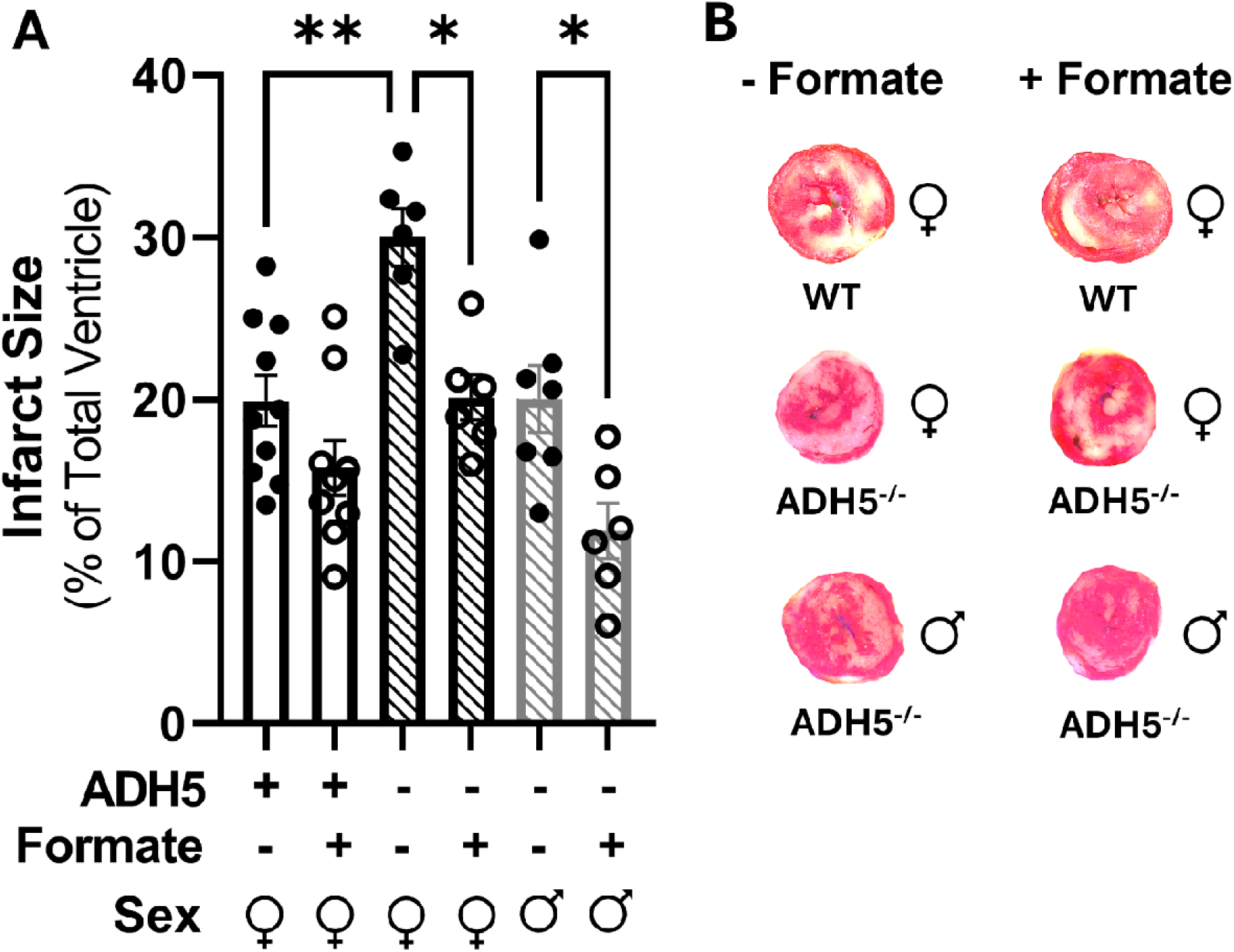
Exogenous formate reduces I/R injury in male and female ADH5^-/-^ hearts. Female WT and female and male ADH5^-/-^ hearts were Langendorff-perfused +/- 30 µmol/L formate and subjected to I/R injury. (A) Infarct size and (B) representative images from female WT and female and male ADH5^-/-^ hearts (**p<0.01 vs. female ADH5^-/-^, *p<0.05 vs. ADH5^-/-^ without formate by ANOVA; n=6-10 hearts/group).

### ADH5 and ALDH2 activity are decreased in OVX WT hearts

Prior work has shown that ADH5 activity is increased in the adult female mouse lung relative to the lungs of juvenile males and females.^26^ Female rat hearts also have increased ALDH2 phosphorylation and activity vs. males, whereas OVX reduces cardiac ALDH2 activity but this activity can be restored with estrogen replacement.^13^ As such, these findings support a role for estrogen in the regulation of ADH5 and ALDH2.

Therefore, we further examined a role for ADH5 and ALDH2 in female-specific cardioprotection by examining the effect of OVX on ADH5 and ALDH2 expression and activity in the mouse heart to determine potential estrogen-dependent effects. Interestingly, we found that despite a significant reduction in ALDH2 protein expression in intact female WT hearts compared to OVX WT hearts (Fig. 2A), ALDH2 activity was significantly increased in intact female WT hearts (Fig. 2B). We also found ADH5 expression to be similar between intact and OVX WT hearts (Fig. 2C), but ADH5 activity was significantly decreased in OVX WT hearts (Fig. 2D). Although ADH5 and ALDH2 activity was reduced in OVX WT hearts, we also found that formaldehyde levels were significantly decreased in OVX WT hearts compared to intact females (Fig. 2E). These findings support a potential role for estrogen in the regulation of ADH5 and ALDH2 expression and activity.

**Figure 2.**
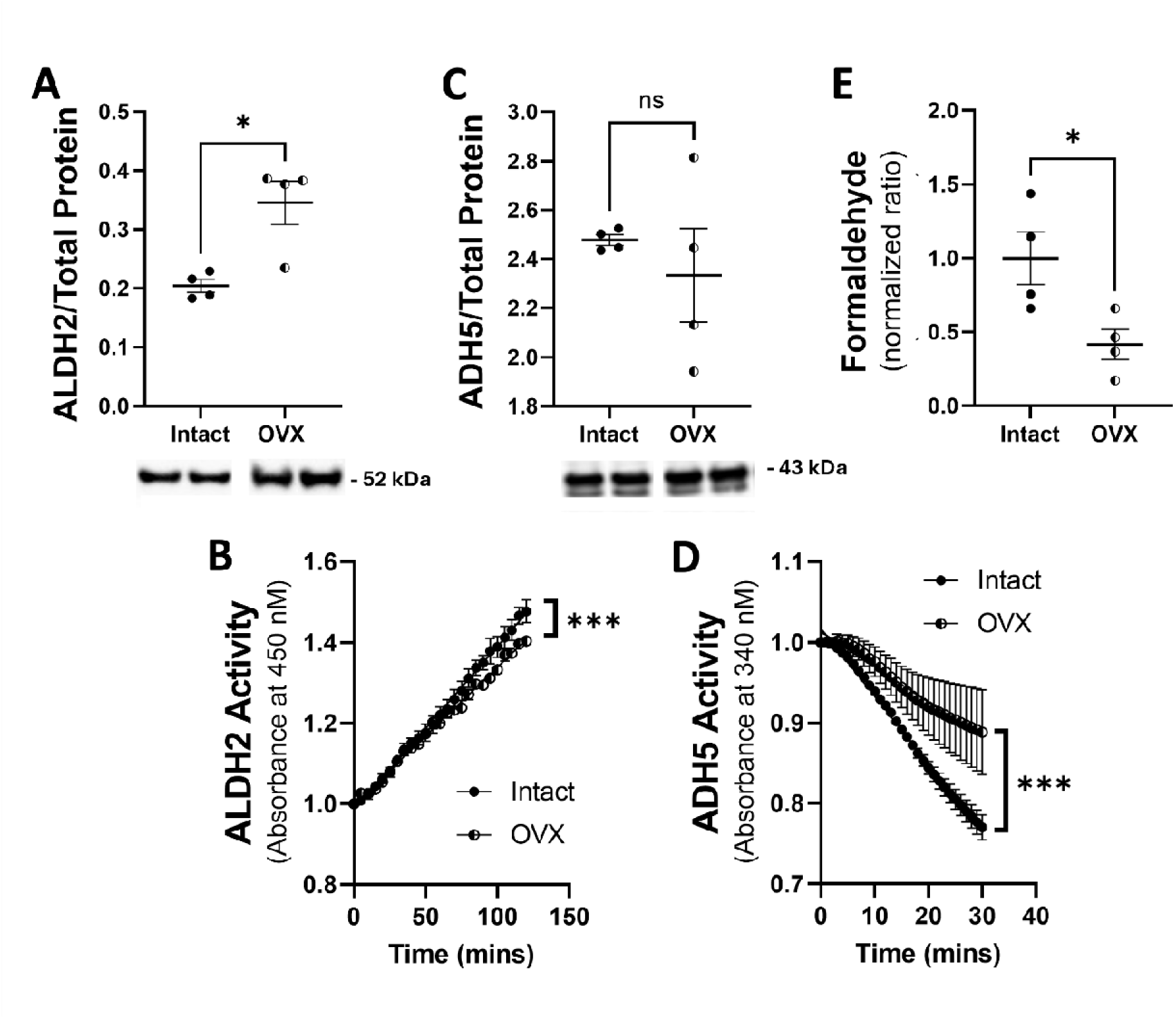
ALDH2 and ADH5 activity, and formaldehyde levels are lower in OVX WT hearts. (A) ALDH2 protein expression via SDS-PAGE and western blot and (B) activity via NADH-linked assay in intact and OVX WT hearts (*p<0.05 by Students t-test, ***p<0.0001 by simple linear regression analysis of slope, n = 4). (C) ADH5 protein expression via SDS-PAGE and western blot and (D) activity via NADH-linked assay in intact and OVX WT hearts (*p<0.05 by Students t-test, ***p<0.0001 by simple linear regression analysis of slope, n = 3-4 hearts/group). (E) Formaldehyde levels were assessed in intact and OVX WT hearts (*p<0.05 by Students t-test, n=4 hearts/group).

### Formate does not rescue OVX WT hearts from ischemic injury

Since ADH5 and ALDH2 activity were both reduced in OVX WT hearts (Fig. 2), we hypothesized that formate production may be similarly impaired. Furthermore, we hypothesized that exogenous formate may provide a rescue for OVX WT hearts subjected to I/R injury. OVX WT hearts were Langendorff-perfused with or without 30 μmol/L formate for 20 minutes, subjected to 25 minutes of ischemia, and then re-perfused with 30 μmol/L formate for 90 minutes. However, formate perfusion did not yield a protective infarct-sparing benefit for OVX WT hearts (Fig. 3). These findings suggest that formate alone is not sufficient to rescue OVX WT hearts and reduce infarct size.

**Figure 3.**
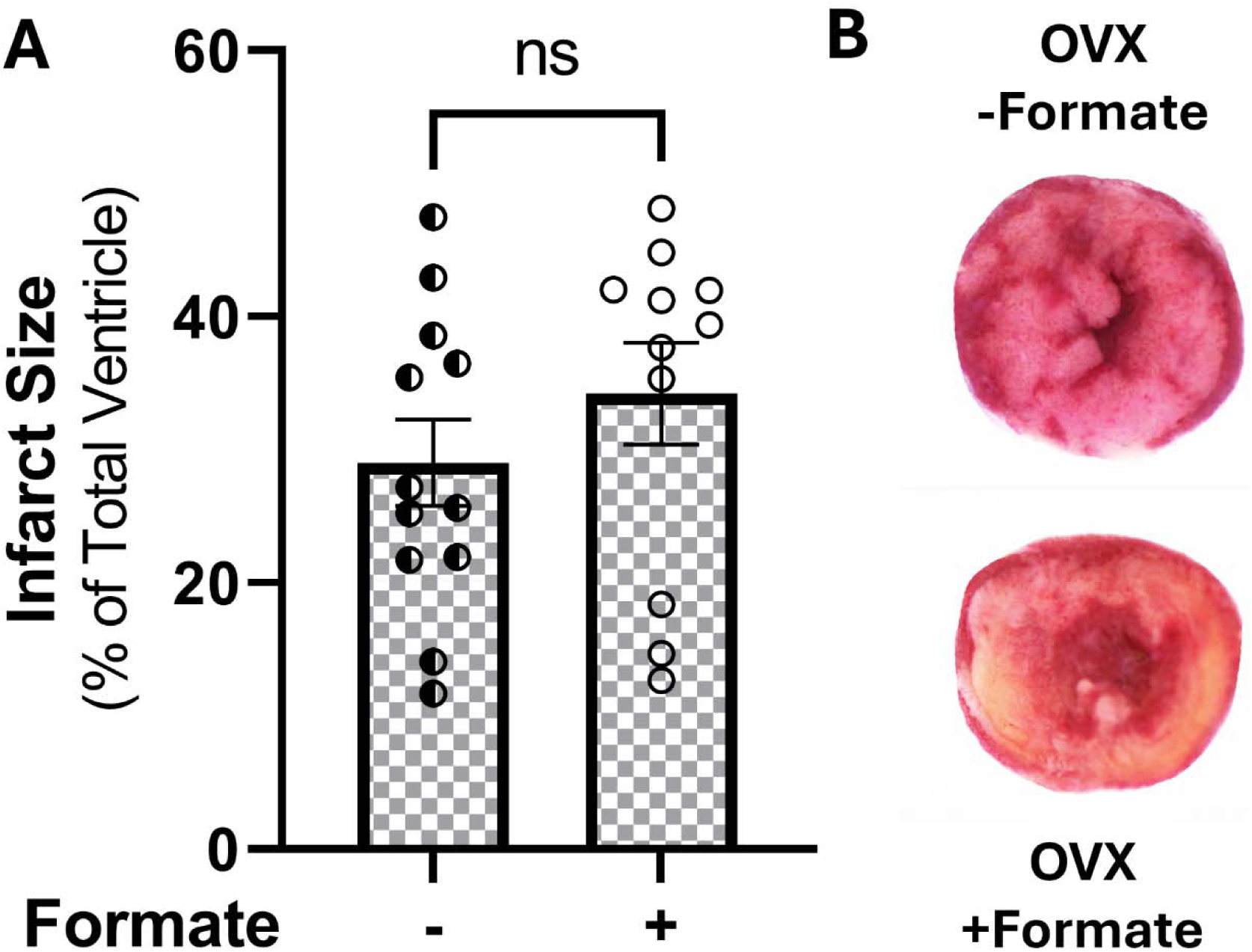
Exogenous formate failed to reduce ischemic injury in OVX WT hearts. OVX WT hearts were Langendorff-perfused +/- 30 µmol/L formate and subjected to I/R injury. (A) Infarct size and (B) representative images from OVX WT hearts (n=11-12 hearts/group).

### Differential expression of one-carbon metabolism enzymes

We next sought to determine a potential mechanism for the lack for formate-induced protection in OVX WT hearts. Since formate is used in OCM and OCM is a major source of formaldehyde generation,^27, 28^ we next explored the differential expression of a panel of OCM enzymes in intact and OVX WT hearts. We found that the protein expression of lysine specific histone demethylase (LSD1) (Fig. 4A) and mitochondrial methylenetetrahydrofolate dehydrogenase (MTHFD2) (Fig. 4B) were both significantly reduced in OVX WT hearts compared to intact females. Conversely, serine hydroxymethyltransferase (SHMT1) (Fig. 4C) protein expression was significantly increased in OVX WT hearts. These findings are notable, as SHMT1 and LSD1 are major sources of formaldehyde production,^29^ while MTHFD2 is critical for importing formate into OCM.^30^ Additional OCM enzymes (methionine synthase, methylene tetrahydrofolate dehydrogenase 1 like [MTHFD1L], methylene tetrahydrofolate reductase [MTHFR], SHMT2, thymidylate synthase) were also evaluated. No significant differences were noted in protein expression (data not shown), however. Together, these findings suggest that estrogen regulates the expression of certain OCM enzymes in the female heart. Moreover, these findings further support a potential mechanism whereby MTHFD2 downregulation may block formate-induced cardioprotection.

**Figure 4.**
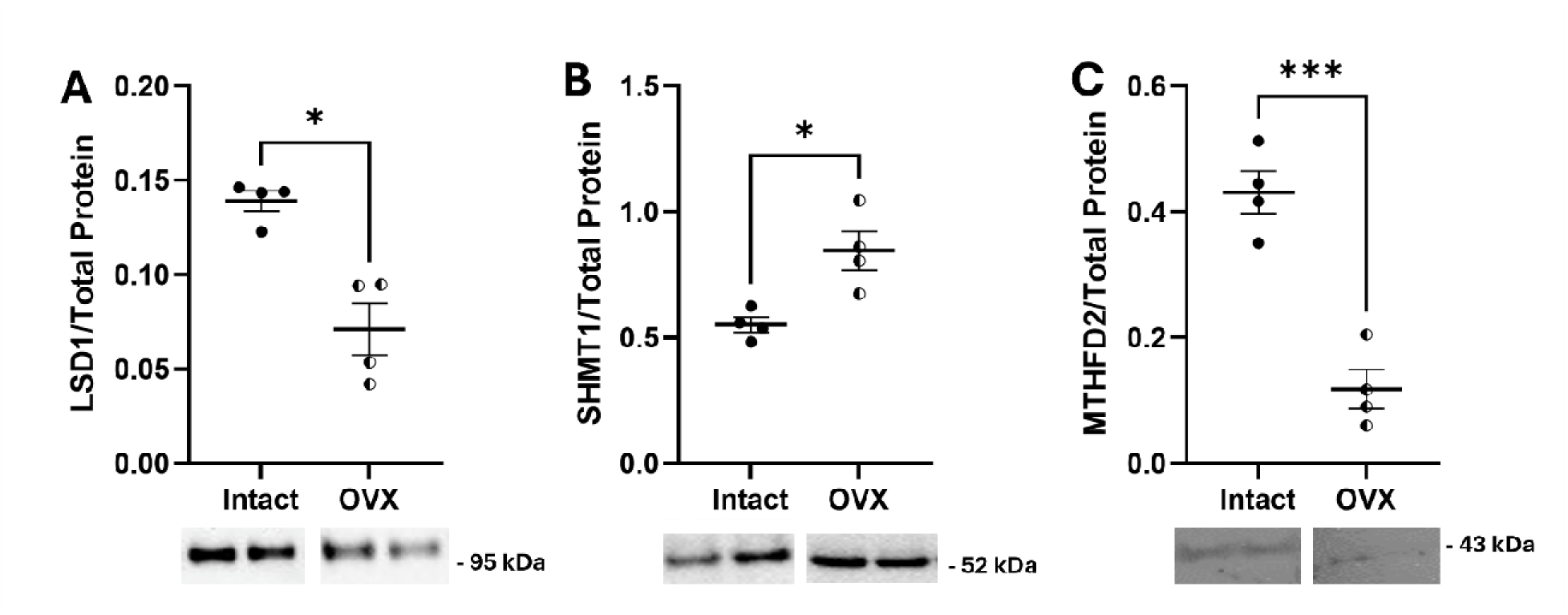
OCM enzyme expression is altered in OVX WT hearts. Protein expression via SDS-PAGE and western blot for (A) LSD1 (*p<0.05 by Students t-test, n=4 hearts/group), (B) SHMT1 (*p<0.05 by Students t-test, n=4 hearts/group) and (C) MTHFD2 (***p<0.001 by Students t-test, n=4 hearts/group).

### Inhibition of MTHFD2 exacerbates I/R injury in intact female hearts

Since MTHFD2 is necessary for importing formate into OCM, we next examined the effect of MTHFD1/2 inhibition on endogenous cardioprotection in intact female WT hearts. Intact female WT hearts were Langendorff-perfused with or without the MTHFD1/2 inhibitor 10 μmol/L LY-345899 (MedChemExpress, Monmouth Junction, NJ) for 20 minutes, subjected to 25 minutes of ischemia, and then re-perfused for 90 minutes. Interestingly, we found that MTHFD1/2 inhibition substantially increased infarct size in intact female hearts (Fig. 5A-B). These findings further support a role for formate and OCM in female-specific cardioprotective signaling.

**Figure 5.**
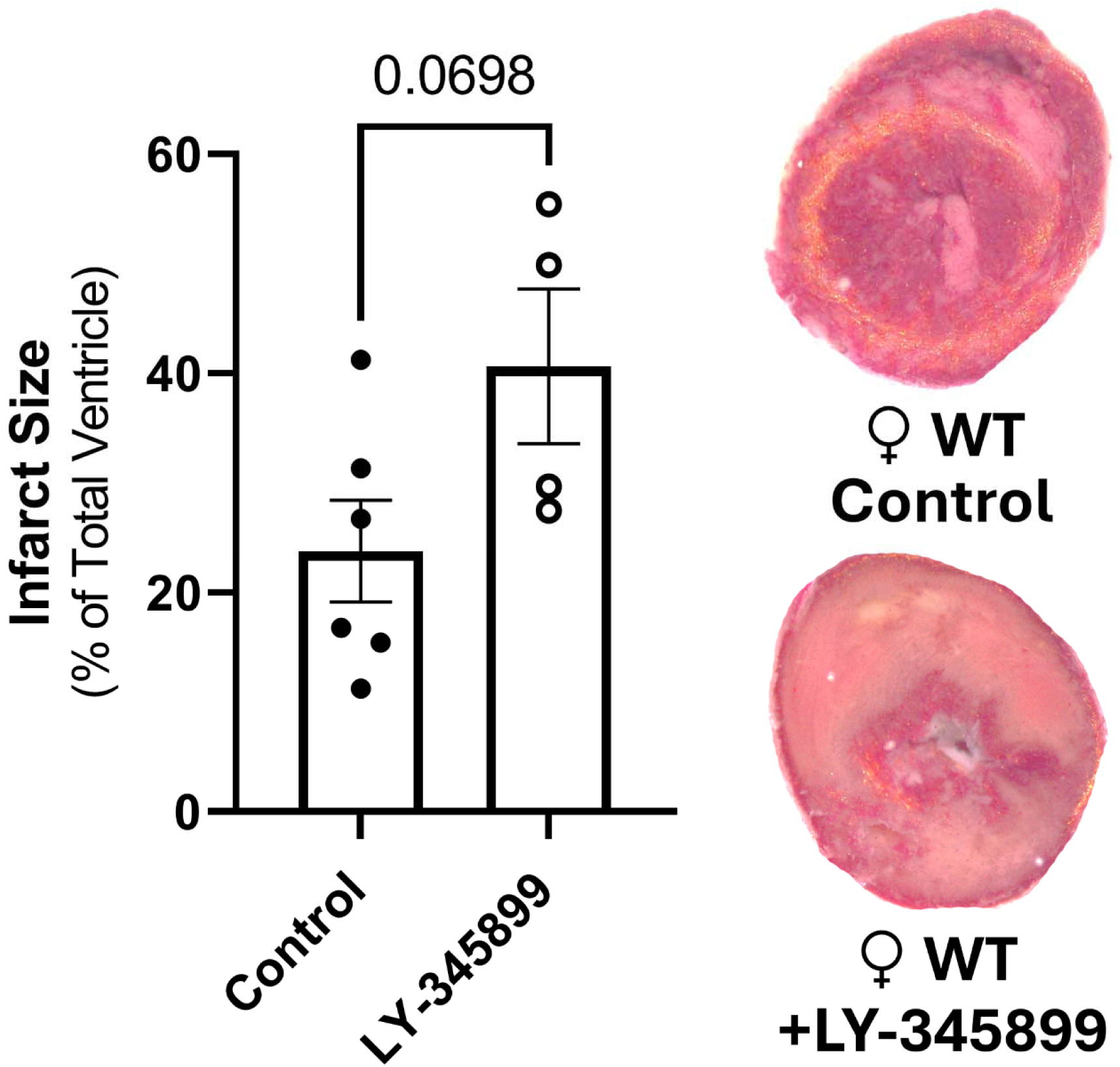
MTHFD1/2 enzyme inhibition blocks sex-specific protection in intact female WT hearts. Intact female WT hearts were Langendorff-perfused +/- 10 µmol/L LY-345899 (MTHFD1/2 inhibitor) and subjected to I/R injury. (A) Infarct size and (B) representative images from intact female WT hearts (p=0.0698 by Students t-test; n=4-6 hearts/group).

## DISCUSSION

Herein, we demonstrate for the first time that formate rescues female ADH5^-/-^ hearts from ischemic injury (Fig. 1). Furthermore, we found that inhibition of MTHFD1/2, which is necessary for importing formate into OCM, blocked female-specific cardioprotection in WT female hearts and exacerbated I/R injury (Fig. 5). Conversely, formate was not effective at reducing infarct size in OVX WT hearts (Fig. 3), despite reduced activity of the formate-generating enzymes ADH5 and ALDH2 in OVX WT hearts (Fig. 2). OVX WT hearts did show a decrease in the expression of MTHFD2 (Fig. 4), which may explain why formate was ineffective at reducing infarct size. Together, our findings support formate and OCM as critical components of sex-dependent cardioprotection, and estrogen as a key regulator of female-specific protective signaling.

We previously showed that ADH5 knockout or inhibition increased post-ischemic formaldehyde levels and exacerbated I/R injury in female hearts.^10^ Interestingly, these effects were reversed upon activation of the formate-generating enzyme ALDH2^10^ or via administration of exogenous formate (Fig. 1). However, exogenous formate failed to provide an infarct-sparring effect in WT female hearts. As such, higher ADH5 activity in WT female hearts vs. male^8, 10^ and OVX WT hearts (Fig. 2) may yield sufficient formate for protective signaling, and thus additional formate may be redundant and not protective. Indeed, recent findings suggest that both ADH5 and ALDH2 generate up to 50% of circulating formate.^29^ Consistent with increased ADH5 and ALDH2 activity in females, intact female mice have higher plasma formate levels compared to males.^35^ Plasma formate is also higher in pre-menopausal women compared to age-matched men,^32^ which may be why formate does not yield additional protection in WT female hearts. Thus, our findings support formate as a critical metabolite for cardioprotective signaling in female hearts, but the underlying protective mechanism(s) remains to be defined.

### Role of formate and OCM

Although formate was discovered as a noxious component of ant and bee venom, it is also produced in mammalian cells, circulating at 20-100 μM in humans and other mammals.^31–34^ Formate has antioxidant properties,^28^ and can be used as a one-carbon donor^29^ to further maintain redox homeostasis for the synthesis of antioxidants including NADPH and glutathione.^30^ Moreover, formate may also contribute to the salvage and recycling of the critical NOS cofactor tetrahydrobiopterin (BH_4_) via OCM.^31^ Importantly, our findings suggest a novel role for OCM in female-specific cardioprotective signaling, as blockage of formate import into OCM via MTHFD1/2 inhibition was sufficient to exacerbate I/R injury in WT female hearts (Fig. 5). Additionally, prior work has shown that the OCM donor folate enhances eNOS function and lessens myocardial ischemic injury by reducing eNOS uncoupling.^32^ Folate treatment of endothelial cells also increases BH_4_ levels and NO production. Formate can enter OCM and be synthesized into 5-methyl-THF by a series of enzymatic reactions,^33^ which can improve eNOS function.^33^ As such, 5-methyl-THF can restore NO by mimicking BH_4_ and enabling eNOS function.^34^ Therefore, formate has the potential to support antioxidant signaling and eNOS function, both of which are enhanced in female hearts, thus combatting redox dysregulation in the heart during I/R injury. However, additional studies are needed to investigate formate production, utilization, and flux within OCM in the female heart to determine underlying protective mechanisms for formate and OCM.

### Role of estrogen in protective signaling

We used an OVX model to evaluate the effect estrogen on ADH5 and ALDH2 activity and found that OVX decreased the activity of both ADH5 and ALDH2 (Fig. 2). As such, our findings suggest that the function of ADH5 and ALDH2 may be regulated by estrogen, which is consistent with prior studies demonstrating that estrogen increases ADH5 activity in the female lung.^26^ OVX rat hearts also have reduced ALDH2 activity.^13^ Furthermore, given the reduction in ADH5 and ALDH2 activity, and formaldehyde levels in OVX WT hearts (Fig. 2), we hypothesized that formate production may be reduced with ovariectomy and examined whether exogenous formate would yield a cardioprotective benefit. However, exogenous formate failed to reduce infarct size in OVX WT hearts (Fig. 4). Since formate is used by OCM and OCM is a source of formaldehyde production, we examined OCM enzyme expression and found lower expression of LSD1 (Fig. 3), which participates in formaldehyde-producing demethylation reactions.^28^ Conversely, expression of SHMT1, which releases formaldehyde by cleaving serine to glycine,^29^ was higher in OVX WT hearts compared to intact females (Fig. 3). As such, expression changes in LSD1 and SHMT1 may be responsible for alterations to myocardial formaldehyde levels, but further confirmatory experiments are necessary. Additionally, MTHFD2 expression was also reduced in OVX WT hearts compared to intact WT females (Fig. 3). Since MTHFD2 is responsible for importing formate into the OCM pathway, it is possible that formate utilization by OCM may be impaired in the OVX WT heart. As such, a multi-pronged approach may be necessary to reduce ischemic injury in OVX WT hearts. While ADH5/ALDH2 activity is reduced in OVX WT hearts, it is possible that this level of activity may still generate a sufficient level of endogenous formate, rendering additional formate non-beneficial. Nonetheless, our findings support estrogen as a critical regulator of the activity and expression of ADH5 and ALDH2, as well as the expression of certain OCM enzymes, but additional research is needed to establish estrogen-dependent links.

### Study Limitations

There are several limitations in this study that require acknowledgement. Firstly, we only evaluated one dose of formate, while physiological levels of circulating formate are fall in the range of 20-100 μmol/L in mammals.^31–34^ As such, the 30 μmol/L formate dose used in our current study may be insufficient to drive protection in intact and OVX WT female hearts. The timing of formate administration may also be critical, as we only administered formate for 20 minutes before ischemia and during the entirety of reperfusion. Therefore, future studies will need to examine the dose- and time-dependent effects of exogenous formate administration. The timing of ovariectomy and the estrogen washout period may also affect the results of our study, so future studies will need to evaluate ovariectomy at different ages and with different washout periods.

## Conclusions

The results of our study support a critical role for ADH5/ADH2, formate, and OCM in female-specific cardioprotective signaling. Specifically, we found that exogenous formate significantly reduces ischemic injury in hearts lacking ADH5. Furthermore, we found that while OVX decreases ADH5 and ALDH2 activity in the heart, exogenous formate does not provide protection from ischemic injury, possibly due to altered OCM enzyme expression. Indeed, inhibition of formate import into the OCM pathway exacerbated ischemic injury in intact female WT hearts. Together, our findings identify formate utilization by OCM as a potentially novel mediator of ischemic injury in female hearts, particularly in the absence of ADH5, but exogenous formate alone does not confer cardioprotection in the context of estrogen depletion. This work represents an important first step in understanding the role of ADH5/ALDH2, formate and OCM in female-specific cardioprotective signaling.

## GRANTS

This work was supported by the National Institutes of Health [T32 ES007141 (HG), F31 HL165820 (HG), R21 HL157800 (MK) and R01 HL136496 (MK)].

## DISCLOSURES

The authors have nothing to disclose.

## Notes

### Competing Interest Statement

The authors have declared no competing interest.

